# Estimating cell cycle model parameters using systems identification

**DOI:** 10.1101/035766

**Authors:** Edwin Juarez, Ahmadreza Ghaffarizadeh, Samuel H. Friedman, Edmond Jonckheere, Paul Macklin

## Abstract

A current challenge in data-driven mathematical modeling of cancer is identifying biologically-relevant parameters of mathematical models from sparse and often noisy experimental data of mixed types. We describe a cell cycle model and outline how to use the Optimization Toolbox in Matlab to estimate its timescale parameters, given flow cytometry and cell viability (synthetic) data, and illustrate the technique with simulated data. This technique can be similarly applied to a variety of cell cycle models, particularly as more laboratories begin to use high-content, quantitative cell screening and imaging platforms. An advanced version of this work (CellPD: cell line phenotype digitizer) will be released as open source in early 2016 at MultiCellDS.org.

## I. INTRODUCTION

Cell cycle time scales are parameters often needed when developing a mathematical model to describe a (cancer) cell population. Measuring these time scales experimentally can be very challenging, and it often requires technologies that are not accessible in most labs. However, using cell flow cytometry [1] we can measure the fraction of cells at three different stages of the cell cycle (G_0_/G_1_, S, G_2_/M). Additionally, propidium iodide (PI) can be used to measure the unviable/apoptotic fraction of the same cell population [2]. Combining these fractions with an automated cell counter (such as the Bio-Rad TC20), we can measure the number of cells at each stage of the cell cycle, and the number of cells undergoing apoptosis.

To study the cell cycle of a cell line, in the lab we would typically record these measurements at a few time points per microenvironmental condition. In this example, we have synthetized data by running the model with a given set of parameters (referred to as the “true parameters”) and multiplying each sample point by a “noise” factor *f* with Gaussian distribution (with a mean of 1 and a standard deviation of 0.2). This process was repeated 3 times (the mean and standard deviation of those 3 samples was computed) to mimic the low number of replicates in many experiments. Fig. 1 summarizes these synthetic data. The methods described in this example can be directly applied to lab data under each specified microenvironmental condition to understand the impact of the microenvironment on the parameters of the model (e.g. cell cycle timescales).

**Figure 1.**
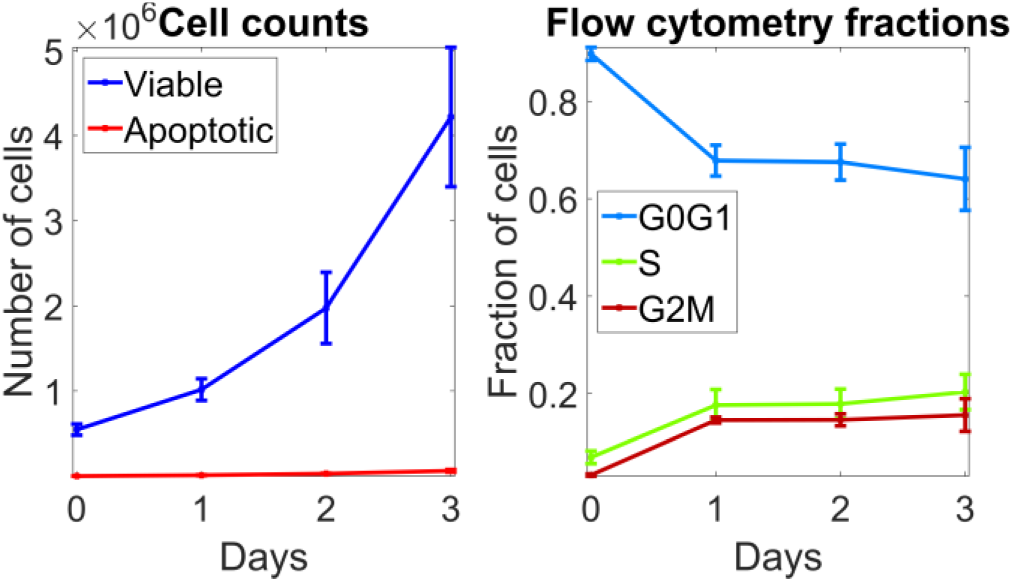
- Synthetic data representative of experimental data. Sample mean ± standard deviation are plotted.

**Authors’ note:** The methods described in this work will be released in a more advanced form through the CellPD (cell line phenotype digitizer) tool. Please check MathCancer.org and MultiCellDS.org/Tools.php for the most up-to-date project information and downloads.

## II. MODEL DESCRIPTION AND PARAMETER ESTIMATION

We can use a system of ordinary differential equations (oDEs) to describe the cell cycle dynamics:

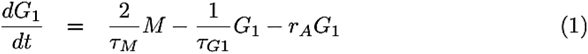

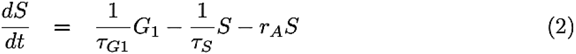

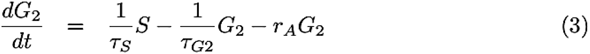

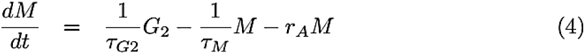

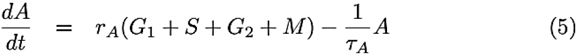

Each equation represents the net change on the number of cells in each modeled stage of the cell cycle (G_0_/G_1_, S, G2, and M) and nonviable (apoptotic) cells (*A*). The parameters (*τ_G_*_1_,*τ_S_*,*τ_G_*_2_,*τ_M_*,*τ_A_*,*r_A_*) represent the mean values of an unknown distribution and are, in general, microenvironment- and time-dependent. For this demonstration, we assume them to be constant in time, and we also assume that the microenvironment does not change significantly enough to modify the value of those parameters during the experiment. Our goal will be to find the set of parameters that best describes the data.

In order to find this optimal parameter set, we select an initial guess for each of the parameters based on literature [3–5] and observations from the lab. We define an error vector, 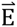, to quantify the discrepancy between the experimental measurements and the model predictions. For any data observation time *t*, we define the raw error vector 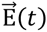 to be the difference between the simulated and experimental values of each cell –sub-population. In order to account for measurement errors, we divide each vector component by its corresponding coefficient of variation (CV); this gives the greatest weighting to data with the least uncertainty. The overall error vector 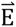 concatenates each sample time’s error vector. In the example presented here, after including cell viability, flow cytometry and cell count data we have 5 “Target” populations: Viable, Apoptosing, G_0_/G_1_, S, and G_2_/M cells. Each of these target populations have 4 time points, so 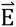 has 20 elements. The sum of the squares of the elements on 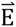 can be computed to find the Sum of Squared Errors (SSE). We can now define an optimization problem by attempting to minimize the SSE while maintaining the timescales within predefined constraints. This minimization is performed by using Matlab’s lsqnonlin function.

Sample code to generate the synthetic data and estimate the timescales of the cell cycle model used in this example will be made available for download from:

http://MathCancer.org/NCI_handbook_parameters, and http://MultiCellDS.org/Tools.php.

## III. RESULTS OF METHOD

After applying the optimization described above, to the optimal parameters (as shown in Table 1, column “Estimated parameter, dataset 1”) lead to a good fit of the data as shown in Fig. 2. But it is worth noting that due to the low number and sparse frequency of the samples, the true parameters could not be recovered exactly. But if we repeat this method and we sample the data more often, more times, and with smaller noise, we can recover the true parameters even if we start from the same initial guess as before. Table 1, column “Estimated parameter, dataset 2” shows the results of the optimization after sampling the data twice per day, and take 30 samples each time with a Gaussian noise with mean 1 and standard deviation of 0.001. Synthetic dataset 2 is virtually impossible to replicate in the lab given the large number of samples and the low level of measurement error required, but it shows that it is possible to extract the true parameters given a sufficiently clean set of data. Furthermore, even with the noisier dataset, we can obtain parameters that not only lead to a good simulation fit but are also close to the true parameters.

**Table 1:**
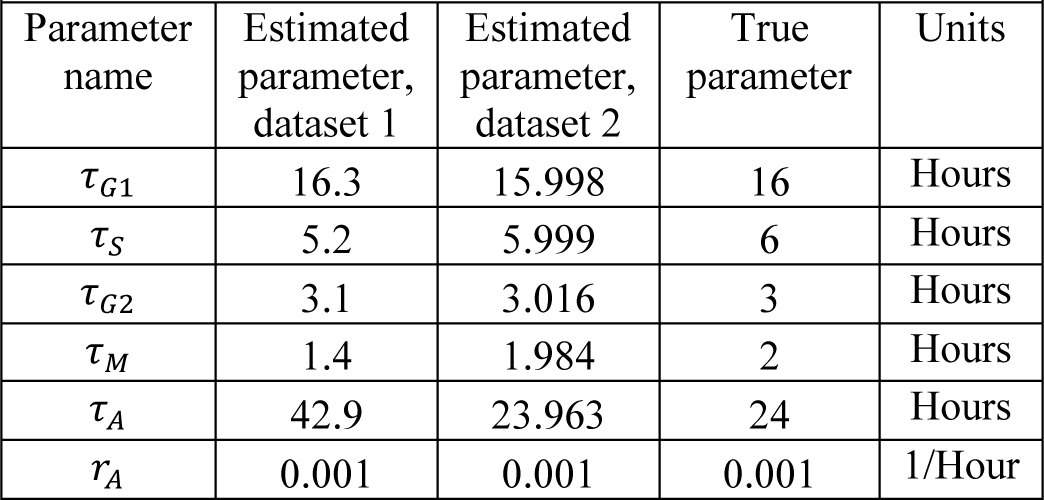
Optimization Results

| Parameter name | Estimated parameter, dataset 1 | Estimated parameter, dataset 2 | True parameter | Units |
| --- | --- | --- | --- | --- |
| $\tau_{G1}$ | 16.3 | 15.998 | 16 | Hours |
| $\tau_S$ | 5.2 | 5.999 | 6 | Hours |
| $\tau_{G2}$ | 3.1 | 3.016 | 3 | Hours |
| $\tau_M$ | 1.4 | 1.984 | 2 | Hours |
| $\tau_A$ | 42.9 | 23.963 | 24 | Hours |
| $r_A$ | 0.001 | 0.001 | 0.001 | 1/Hour |

**Figure 2:**
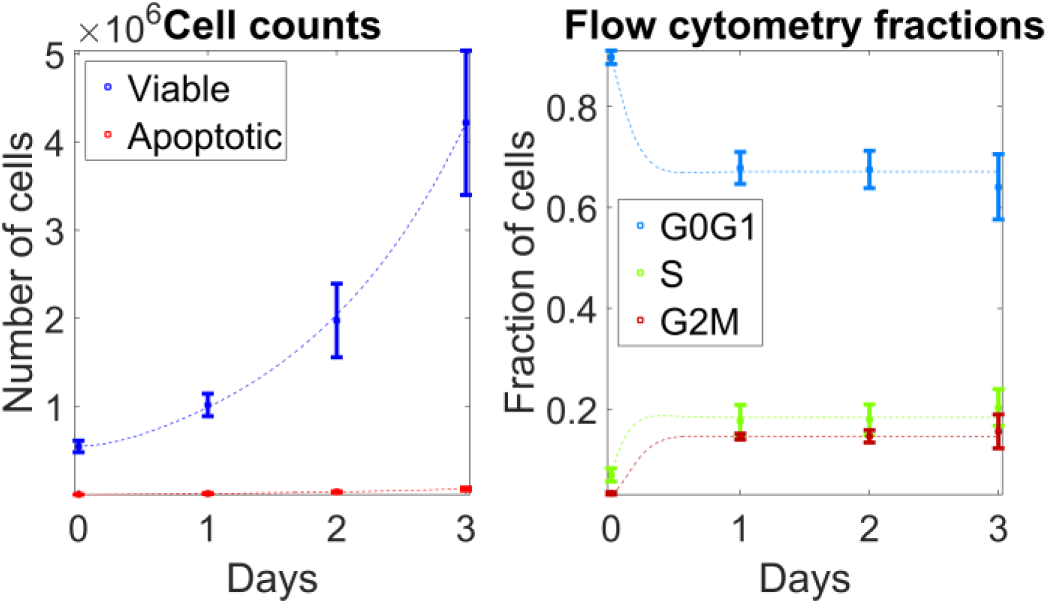
Simulation results using estimated parameters from dataset 1 (3 samples every 24h with a Gaussian noise of mean 1 and standard deviation of 0.2). Sample mean ± standard deviation are plotted, dotted lines represent simulation output.

## IV. TYPE OF SETTINGS IN WHICH THESE METHODS ARE USEFUL

This method can be applied in virtually the same way to other cell cycle models, where the errors and model are chosen to suit the available data. The same methodology can be used to calibrate agent-based models by defining an error metric and tuning simulation parameters to minimize that error function. Lastly, we note that the optimization technique can be iterated to help quantify uncertainty in the parameters.

